# Charged Gram-positive species sequester and decrease the potency of pediocin PA-1 in mixed microbial settings

**DOI:** 10.1101/2020.04.13.039628

**Authors:** Vikas D. Trivedi, Nikhil U. Nair

## Abstract

Antimicrobial peptides (AMPs) have gained attention recently due to increasing antibiotic resistance amongst pathogens. Most AMPs are cationic in nature and their preliminary interactions with the negatively charged cell surface is mediated by electrostatic attraction. This is followed by pore formation, which is either receptor-dependent or -independent and leads to cell death. Typically, AMPs are characterized by their killing activity using bioactivity assays to determine host range and degree of killing. However, cell surface binding is independent from killing. Most of the studies performed to-date have attempted to quantify the peptide binding using artificial membranes. Here, we use the narrow-spectrum class IIa bacteriocin AMP pediocin PA-1 conjugated to a fluorescent dye as a probe to monitor cell surface binding. We developed a flow cytometry-based assay to quantify the strength of binding in target and non-target species. Through our binding assays, we found a strong positive correlation between cell surface charge and pediocin PA-1 binding. Interestingly, we also found inverse correlation between zeta potential and pediocin PA-1 binding, the correlation coefficient for which improved when only Gram-positives were considered. We also show the effect of the presence of protein, salt, polycationic species, and other non-target species on the binding of pediocin PA-1 to the target organism. We conclude that the of presence of highly charged non-target species, as well as solutes, can decrease the binding, and the apparent potency, of pediocin PA-1. Thus, these outcomes are highly significant to the use of pediocin PA-1 and related AMPs in mixed microbial settings such as those found in the gut microbiota.

## Introduction

Antimicrobial peptides (AMPs) that serve multiple functions are produced by both bacteria and higher organisms. AMPs are part of innate immune response in higher organism, whereas in bacteria, they provide growth advantage competitive ecological niches (1). While traditional antibiotics tend to be broad in their spectrum of activities, many bacteriocins target species that are closely-related to the producer organism—often including only a few species and strains (2). Additionally, bacteria are less likely to become resistant to AMPs (3). Also, resistance to conventional antibiotics is also associated with widespread collateral susceptibility to antimicrobial peptides (4). Due to this, AMPs serve as possible solution to the proliferation of antibiotic-resistance (4, 5). AMPs produced by bacteria are often included among ribosomally-synthesized bacteriocins, classified as class I and II based on extent of post-translational modification and range in size from ~2 to 10 kDa. Many are resistant to boiling and extreme pH, often exhibiting extremely potent bactericidal activity at nanomolar concentration (6, 7). Bacteriocins particularly produced by lactic acid bacteria (LAB) are of special importance to food industry due to the GRAS status of the producer strains. Along with being non-toxic to eukaryotic cells, LAB-derived bacteriocins are sensitive to gut proteases such as trypsin, chymotrypsin, and pancreatin complex; thus, they are predicted to have minimum impact on the gut microbiota (8). Moreover, since they are ribosomally-synthesized, they can be engineering through traditional protein and peptide engineering techniques (9). Class II bacteriocins are further divided into 5 subclasses (a, b, c, d, and e), among which pediocin-like bacteriocins (class IIa), contain a conserved N-terminal YGNGV domain and are one of the most well-studied peptides, with particular interest in food and biotechnological applications due to their innate anti-Listerial activity (10–13). Numerous peptides identified in this subclass have been found to be cationic, containing 37 − 48 amino acid residues, contains one or two disulfide linkages and kill target cells, including the foodborne pathogen *Listeria monocytogenes*, by permeabilizing the cell membrane (14). Pediocin-like bacteriocins exhibit extensive sequence similarity but differ markedly in their target cell specificity (15). Their inhibitory spectrum includes several other genera of gram-positive bacteria such as *Enterococcus*, *Lactobacillus*, *Leuconostoc*, *Pediococcus*, and *Clostridium* and are inactive toward Gram-negative bacteria. These peptides consist of two distinct domains − the N-terminal domain is structured and cationic whereas the C-terminus is unstructured in solution but switches to an amphipathic α-helix structure that terminates with a loop or hairpin-like tail structure upon target binding (16). The mannose phosphotransferase system (Man-PTS) of several Gram-positive bacteria has been confirmed as the specific receptor for pediocin-like bacteriocins (17). It is suggested that receptor-mediated mechanism is most likely behind the specificity of this class of AMPs (18). The initial binding of AMPs to the target cell surface is presumed to be mediated by electrostatic interactions with the positively charged N-terminal domain (19). This is followed by more specific interactions by the C-terminal domain with the Man-PTS, pore-formation, loss of transmembrane potential, and death (20–22). This bactericidal activity can be readily quantified using agar-well diffusion or microtiter plate-based MIC assays. While such assays are useful to identify targets, it gives little to no information on the binding characteristic of the AMPs. Multiple, but indirect approaches have been used to quantitate the binding component of the AMPs to cell surface (7, 19). But these studies have either limited to characterizing the role of charged residues in binding the target species using pull-down assays (19) or using phospholipid vesicles (7). To the best of our knowledge, no study has been performed to quantify the binding of the AMPs using whole cells and determine the binding strength of peptide to target as well as non-target species.

In this study, we first demonstrate that fluorophore conjugates can be used to study pediocin PA-1 behavior and function. Rather than using lipid vesicles, we used live cells to study the binding events. We demonstrate the utility of the conjugate by quantifying the binding constants of pediocin PA-1 with target and a range of non-target bacteria. Further, by using poly-L-lysine, a non-bactericidal polycationic molecule, and zeta potential measurements, we correlate the binding of pediocin to the surface charge and zeta potential. We also show the effect of the presence of protein, salt, poly-cationic species and other non-target species on the binding of pediocin PA-1 to the target organism. We conclude that that the negative charge of Gram-positive bacterial cell surface is directly and strongly correlated with of initial pediocin PA-1 binding and that presence of highly-charged, non-target species, as well as solutes can decrease the binding, and thus the potency of pediocin PA-1. Thus, these outcomes are highly significant to the use of pediocin PA-1 and related AMPs in mixed microbial settings such as those found in the gut microbiota.

## Results and Discussion

### The bioactivity of *Pediococcus acidilactici* UL5 secreted factor(s) is consistent with that of pediocin PA-1

We tested the antimicrobial spectrum of the pediocin PA-1 producer strain, *Pediococcus acidilactici* UL5, using agar-well diffusion assays against few representative members of lactic acid bacteria (LAB) and Gram-negative, *E. coli* DH5α as a control (Fig. 1). The effectiveness of the spend culture medium was evaluated by its ability to inhibit growth of the indicator strains as measured by zone of inhibition (ZOI). Pediocin PA-1 is known to displayed narrow spectrum only against a few species (Table S1) that includes *Lactobacillus coryniformis* B-4390. We found that the bioactivity of spent *P. acidilactici* UL5 supernatant was consistent with that of pediocin PA-1 since it only inhibited *L. coryniformis* (ZOI ≈ 8 mm) and no other members of LAB (Fig. 1a). Therefore, we conclude that the primary antimicrobial secreted is pediocin PA-1.

**Figure 1:**
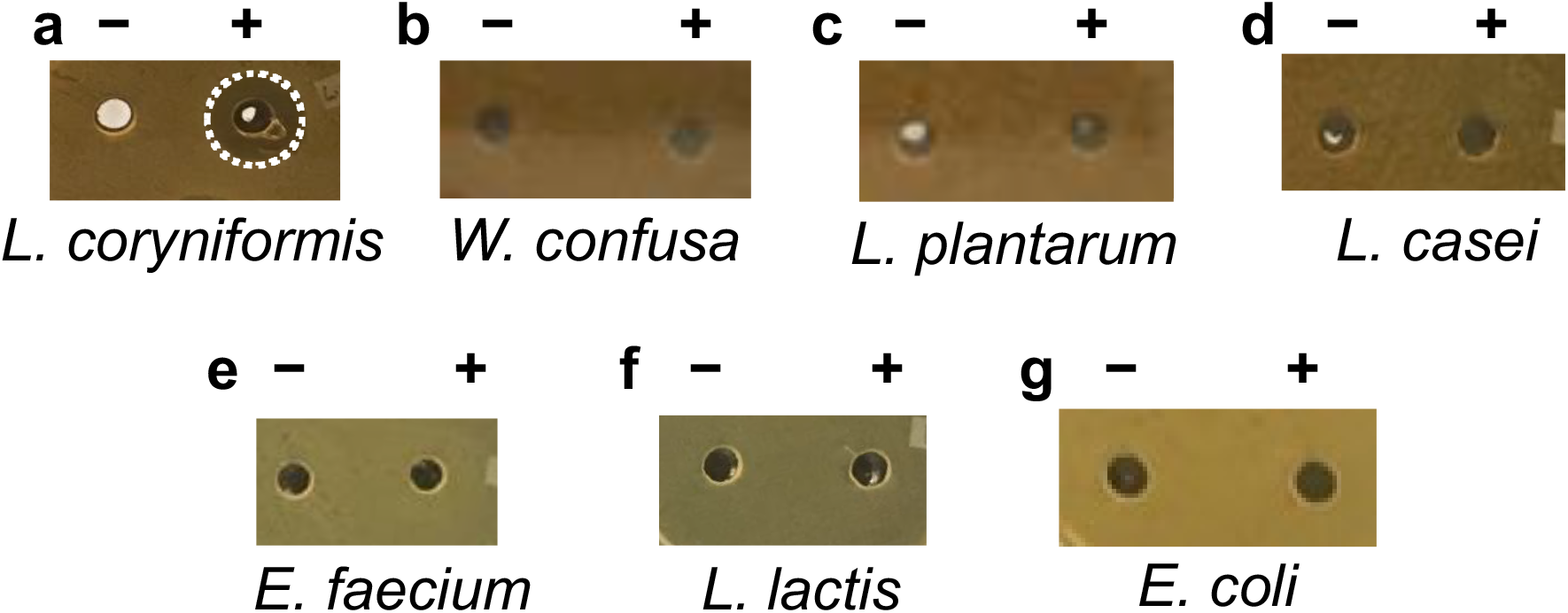
Bioactivity spectrum of *P. acidilactici* UL5 spent supernatant. Supernatant (10 μL) from *Pediococcus acidilactici* UL-5 was used as source of pediocin PA-1 to assess bioactivity against (a) *Lactobacillus coryniformis* B-4390, (b) *Weisella confusa*, (c) *Lactobacillus plantarum* WCSF1, (d) *Lactobacillus casei* B-1922, (e) *Enterococcus faecium* NRRL B-2354, (f) *Lactococcus lactis* MG1363, and (g) *E. coli* DH5α. Only *L. coryniformis* was found to be sensitive. – control; + supernatant; circle highlights the presence of zone of inhibition (ZOI).

### Pediocin PA-1 can be purified by cellular absorption/desorption method

Cell adsorption-desorption has been demonstrated previously as a viable technique to purify bacteriocins from spent media (23). We found that using the producer strain *P. acidilactici* UL5 for adsorption-based purification instead of the target strain, *L. coryniformis*, was more convenient as it minimized the processing steps and hence reduced loss in final yield. Desorption attempted at pH 1.0 and 9.0 yielded similar peptide recovery, but the peptide recovered at the more acidic pH was found to be more stable, which is consistent with data on pediocin PA-1 (23). The recovered peptide, post-dialysis, was lyophilized for long term storage to minimize any loss in activity. The peptide using this method was purified to 189-fold with specific activity of 9,434 AU·mg^−1^ as measured by agar-well diffusion method. Though the molecular mass of pediocin is expected to be 4.5 kDa, we observed the partially purified pediocin to migrate with an apparent mass of 12.5 kDa (Fig. 2a). This is likely due to its highly cationic nature, causing it to migrate slower due to low *m/z* (subsequent analysis by LC-MS enabled a more accurate and precise size determination, *vide infra*). The purified pediocin PA-1, separated on denaturing but non-reducing SDS-PAGE (since this bacteriocin contains two crucial disulfide bonds (24)), was subjected to agar-overlay assay using *L. seeligeri* as indicator organism (Fig. 2b). A single zone of clearing was observed indicating that the purified peptide was indeed pediocin PA-1.

**Figure 2.**
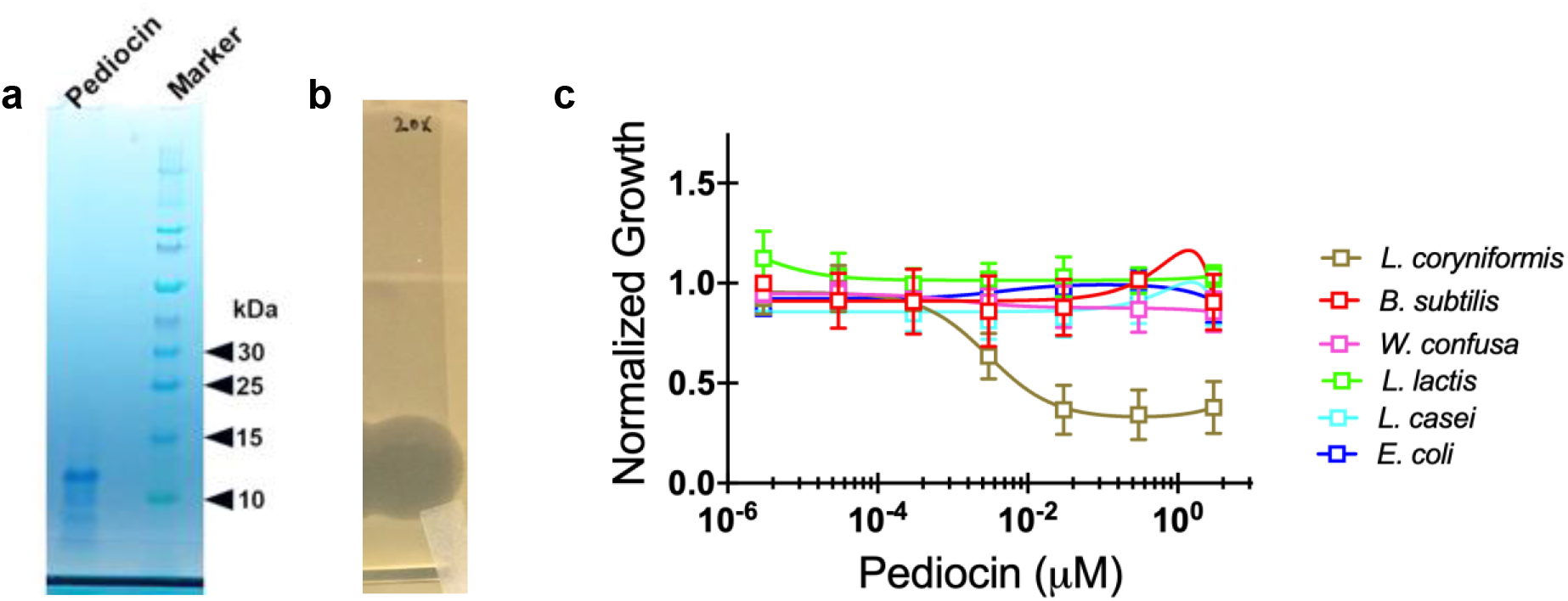
SDS-PAGE analysis and bioactivity of purified pediocin PA-1. (a) Non-reducing SDS-PAGE profile of purified pediocin PA-1 using cell adsorption-desorption method; (b) Agar-overlay assay of purified pediocin using *Listeria seeligeri* B-37019 as an indicator organism. The peptide was subjected to non-reducing SDS-PAGE before performing the assay. (c) Kill-curve of pediocin against the target and the non-target species. EC_50_ for *L. coryniformis* is 3 nM, all other strains are resistant.

In order to determine the potency of the purified pediocin, we generated kill-curves against a range of target microorganisms (Fig. 2c). The concentration range tested was 3 pM to 3 μM following 10-fold serial dilution. As expected, pediocin displayed activity only against *L. coryniformis* with EC_50_ of 3 nM. This is in approximate agreement with published potency pf pediocin PA-1 and related bacteriocins (25–27).

### Fluorescently labelled pediocin PA-1 can be used to detect and quantify binding events

Purified pediocin was conjugated to Atto488 or Atto647 using EDC−NHS (1-ethyl-3-(3-dimethylaminopropyl)carbodiimide − *N*-hydroxysuccinimide) chemistry (Fig. 3a). Atto488 was chosen due to its high fluorescence yield, high photostability, minimum aggregation, and with net charge of −1 it does not significantly impact the overall cationic nature of pediocin. The Atto488-conjugated pediocin was dialyzed and the excess dye was removed using acrylamide desalting column (1 kDa MWCO, Fig. 3b). Pediocin and its fluorescent conjugates Atto488 and 647 were analyzed using MALDI-MS to check for the increase in molecular mass after derivatization (Fig. S1). The partially purified pediocin PA-1 gave a predominant peak at 4,670 Da, which is close to its expected molecular mass. The Atto647 conjugate gave a predominant peak at 5,589 Da, this difference in molecular mass of 919 Da is close to the molecular mass of Atto647-NHS ester (811 Da), suggesting incorporation of single molecule of dye per peptide. We could not detect Atto488 conjugate in MALDI despite repeated attempt or subjecting the peptide conjugate to clean-up using Zip-Tip method. This may be presumably due to poor peptide flight after conjugation. Since conjugation or labeling of the antimicrobial peptide to the fluorescent probes have been reported to alter the antimicrobial activity in some cases (28, 29), pediocin-Atto488 conjugate was also evaluated for its antimicrobial activity (Fig. S2). The antimicrobial results showed that pediocin maintained its antimicrobial activity after conjugation to Atto488/647. However, the results showed that the EC50 was 10-fold lower for both Atto488 and 647 conjugates relative to the unmodified peptide. It is interesting to note that Atto488 has a net negative charge of −1, whereas Atto647 is zwitterionic, but both the conjugates display similar antimicrobial profile suggesting that charge of the dye is not aberrantly altering behavior of the peptide. The conjugate was also tested for its ability to bind *L. coryniformis* using fluorescence microscopy (Fig. S3). Indeed, we could observe the fluorescence signal lining the cell periphery, presumably due to binding of pediocin-Atto488 conjugate on to the cell membrane.

**Figure 3.**
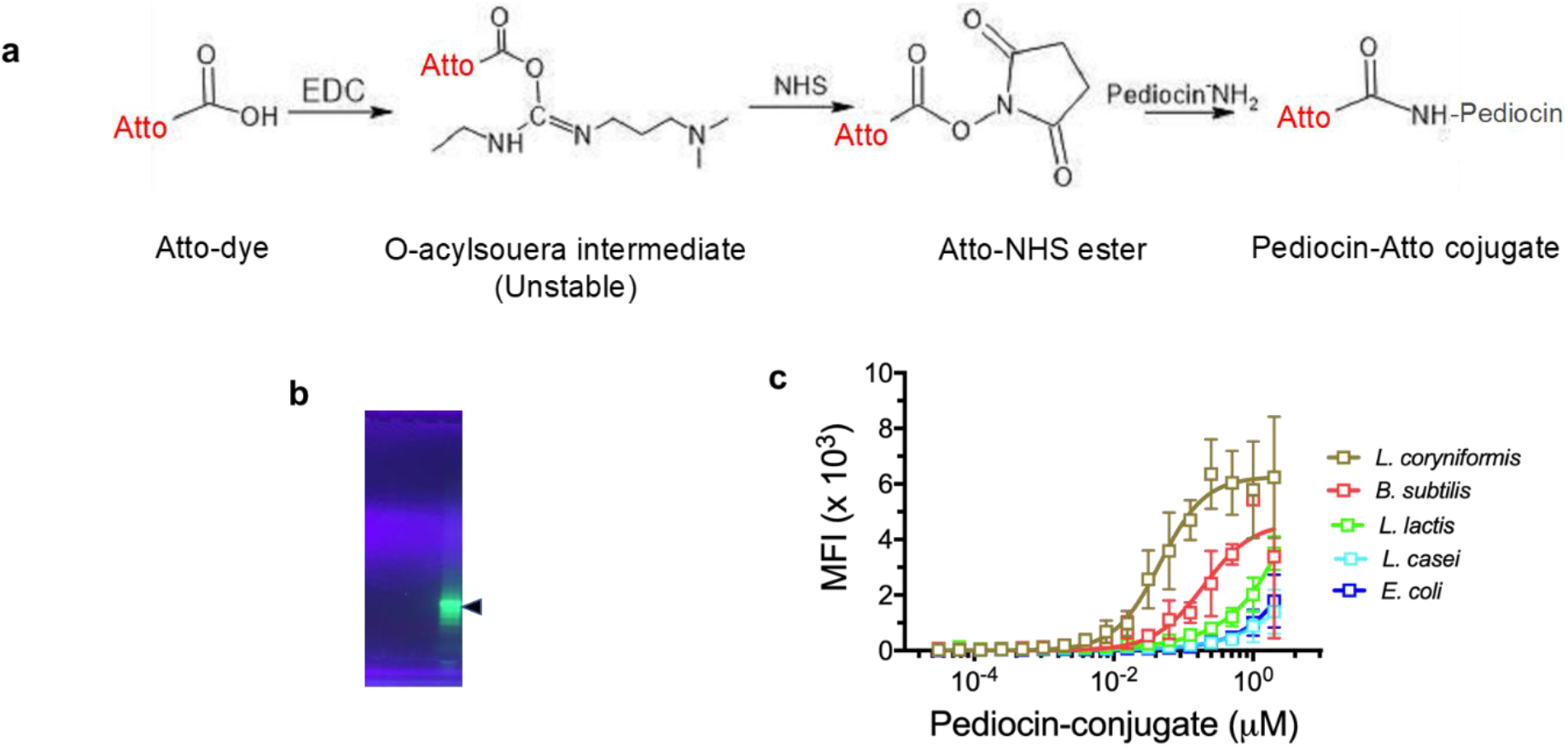
Binding affinity and specificity of pediocin PA-1-dye conjugate. (a) Reaction scheme for EDC/NHS chemistry to conjugate pediocin PA-1 to Atto488/647 dye. (b) UV-fluorescence of pediocin-Atto488 (Ped-488) conjugate after separation on non-reducing SDS-PAGE. (c) Cell-binding curve of Ped-488 against the target and the non-target species. *E. coli* was used as a negative control. *L. coryniformis* demonstrated highest affinity for pediocin with K_D_ = 46 nM.

We developed a flow cytometry-based assay to assess the specificity of pediocin binding to different microorganisms. In order to evaluate this, we initially performed tests to determine the cell density at which to perform the binding experiment (Fig. S4). We evaluated twofold serial dilutions of *L. coryniformis* ranging from OD_600_ 0.05 to 2.0 at a fixed concentration of Ped-488 (0.1 μM). The median fluorescence intensity (MFI) gradually increased with decreasing cell density and approached saturation at OD_600_ ≤ 0.05. Based on this, the binding specificity for all the strains tested was evaluated at OD_600_ of 0.001. This was to ensure that the Ped-488 conjugate was always in molar excess to the binding sites available on the cell surface.

Among the strains tested, as expected *L. coryniformis* showed the highest affinity and maximum median fluorescence for the conjugate (*K*_*D*_ = 46 nM, 6300 MFI) (Fig. 3c, Table 1). The binding curve for the other tested microorganism never attained saturation in the concentration range tested (up to 2 μM). *Bacillus subtilis* showed comparable binding for Ped-488 as indicated by the maximum MFI attained, albeit with lower affinity (*K*_*D*_ = 467 nM). The other strains showed overall weak binding for Ped-488. It is speculated that AMPs like pediocin PA-1 exert their bioactivity through a multistep process consisting of binding, insertion, and pore formation (30). This first step of bacteriocin binding to the membrane is thought to be mediated by electrostatic interaction through the patches of positively charged amino acid residue (7, 19). These residues may permit interaction with phospholipid head groups. This may explain the significant amount of binding displayed by non-target organisms like *B. subtilis*. However, there is substantial evidence in the literature that there is need for a specific receptor like Man-PTS with cognate residues for exertion of bactericidal activity (12, 18, 31).

**Table 1.**
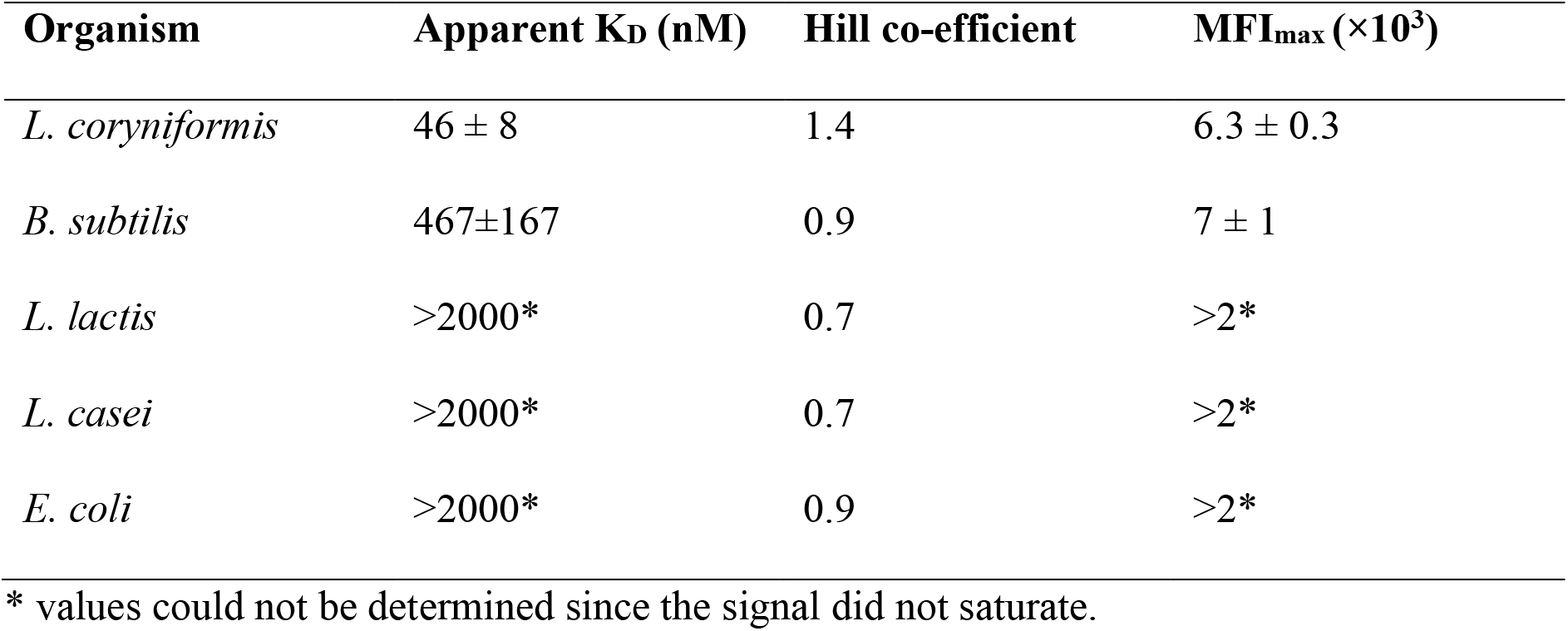
Pediocin PA-1 binding constant to various strains.

### Electrostatic interactions control pediocin PA-1 binding to Gram-positive cell surfaces

As pediocin PA-1 is a cationic antimicrobial peptide, we postulated whether the strength or magnitude of binding has any correlation with cell-surface charge. To test this, we assayed eight different microorganisms for their ability to bind to Ped-488, poly-L-lysine-FITC (PLL-FITC at 0.5 μM) and also measured their zeta potential. *Listeria seeligeri*, which is another organism targeted by pediocin PA-1, showed maximum MFI for both Ped-488 and PLL-FITC (Fig. 4a). We tested the correlation between pediocin and PLL binding using linear regression and Spearman’s correlation coefficient and found positive correlation (r^2^ = 0.84, r_s_ = 0.95, P < 0.001) between pediocin and PLL binding (Fig. 4a). Increase in PLL-FITC binding has previously also been correlated with increased susceptibility to the action of antimicrobial peptides (5, 32, 33). In contrast, zeta potential values did not show a strong correlation with Ped-488 (r^2^ = 0.06, r_s_ = −0.62, Fig. 4b) or PLL-FITC binding (r^2^ = 0.15, r_s_ = −0.62, Fig. 4c). Zeta potential measures the electrochemical property of the cell surface which in turn also reflects surface charge which originates from the negative lipids and lipopolysaccharide (LPS), teichoic acids, or peptidoglycan etc. present in the outer membrane of bacteria. Thus, cells with higher negative zeta potential would be expected to have higher Ped-488 or PLL binding. In our study we did not find this to be true for all the Gram-positive and negative strains tested (Fig. 4b and 4c). If *E. coli* is excluded from the analysis, there appears to be a weak negative correlation between zeta potential and Ped-488 (r^2^ = 0.57, r_s_ = −0.96, P < 0.002, Fig. 4b) or PLL binding (r^2^ = 0.63, r_s_ = −0.96, P < 0.002, Fig. 4c). This may have to do with difference in the cell surface architecture between the Gram-positive and -negative bacterial cells. While changes in cellular zeta potential has been observed upon AMP binding, to the best of our knowledge, no study has correlated zeta-potential with and binding strength with cationic peptides (34).

**Figure 4.**
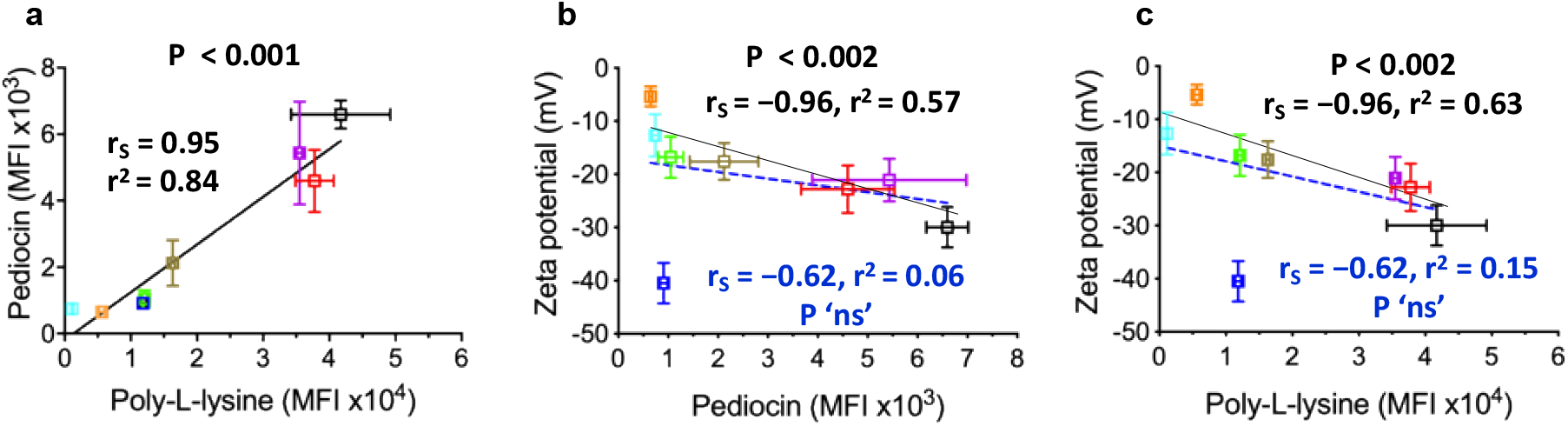
Binding interactions between pediocin PA-1 and poly-L-lysine with cells. Correlation between (a) pediocin PA-1 and poly-L-lysine; (b) pediocin PA-1 and zeta potential; (c) poly-L-lysine and zeta potential for *L. coryniformis* 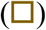, *L. seeligeri* 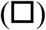, *B. subtilis* 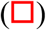, *E. faecium* 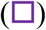, *L. lactis* 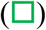, *L. plantarum* 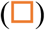, *L. casei* 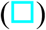. The fit was generated using linear regression (r^2^), the data was analyzed using Spearman correlation model (r_S_). The coefficient for the respective model is indicated along with the P value. The median fluorescence intensity for pediocin and poly-L-lysine was determined with 0.5 μM peptide. In panel b and c, blue dotted line is fit for data including all strains and black solid line is fit for data excluding *E. coli.* ns, non-significant

### Presence of solutes strongly influences pediocin PA-1 binding

In addition, we tested how presence of salt, another protein or charged peptide, or non-target microorganisms affect the binding of pediocin to target species *L. coryniformis*. To begin, the binding of Ped-488 to *L. coryniformis* was tested in presence and absence of NaCl, BSA, and other charged species like poly-D-lysine (Fig. 5a–e). In absence of any additive, *L. coryniformis* displayed significant binding to Ped-488 (Fig. 5b). In the presence of BSA (1 %) and NaCl (100 mM), although 100 % of cells bound to pediocin, the MFI dropped by nearly 10-fold indicating that the solutes disrupted the interactions between the cell surface and the peptide (Fig. 5c). This observation is consistent with earlier studies that have shown an increase in the MIC values for sakacin P (a pediocin PA-1-like class IIa bacteriocin) by 2- to 100-fold in presence of NaCl or MgCl_2_ (7, 35, 36). The study cited above, and our data suggests reduction in the binding of the peptides to target cells due to increased ionic strength because of salts. The presence of poly-D-lysine (PDL, 100 μM) however, displayed significant competition for the peptide binding as 50 % of cells showed no binding at all (Fig. 5d). It was surprising to see the peaks broaden – with cell binding ranging from being unaffected to showing no binding at all. Interestingly, presence of all the additives significantly impacted the binding of the peptide to the target strain *L. coryniformis* as only 7 % of cells showed any significant binding (Fig. 5e). Next, we evaluated the effect of these solutes on binding of Ped-488 to non-target species that lack the cognate receptor/binding site for pediocin PA-1 (*L. casei*, *L. plantarum*, *W. confusa*, *E. faecium*, *B. subtilis* and *E. coli*) in a mixed community setting. The non-target species showed subtle differences in pediocin binding in presence of the additives (Fig. 5f–j). In absence of any additives, 95 % the cells in mixed community showed any significant binding to Ped-488 (Fig. 5g). Compared to *L. coryniformis*, MFI of the Ped-488 bound mixed community was ~3-fold lower, indicating poor avidity. In the presence of BSA and NaCl, only 42 % of the cells showed any significant binding to Ped-488 (Fig. 5h). Also, the residual MFI dropped significantly (5 %). Surprisingly, PDL only dropped the MFI ~2-fold without affecting the percent (%) of cells bound, indicating loss of pediocin binding sites to PDL (Fig. 5i). This contrasts with what we observed for the target species, *L. coryniformis* in which half the population showed no Ped-488 binding. The presence of all the additives did not significantly impact the fraction of population showing Ped-488 binding (Fig. 5j). These results appear counterintuitive at first, but presence of salt not only interferes with the electrostatic interaction between the peptide and the cell but also affects the hydrophobicity of the cell membrane (37). And due to this multifaced effect of salts, it may impact the binding of pediocin to the non-target species for the lack of cognate receptor. Also, the mixed community sample contains cells with varying level of surface charge and lipid composition compared to single population of *L. coryniformis*, which would explain why PDL did not significantly alter their pediocin interaction. In addition, *L. coryniformis* displayed all or none binding in presence of highly charged molecule like PDL suggesting cooperativity in pediocin binding to the target species. This is further supported by the mildly cooperative Hill-coefficient (*n* = 1.4) obtained from cell binding experiments for *L. coryniformis* (Table 1).

**Figure 5.**
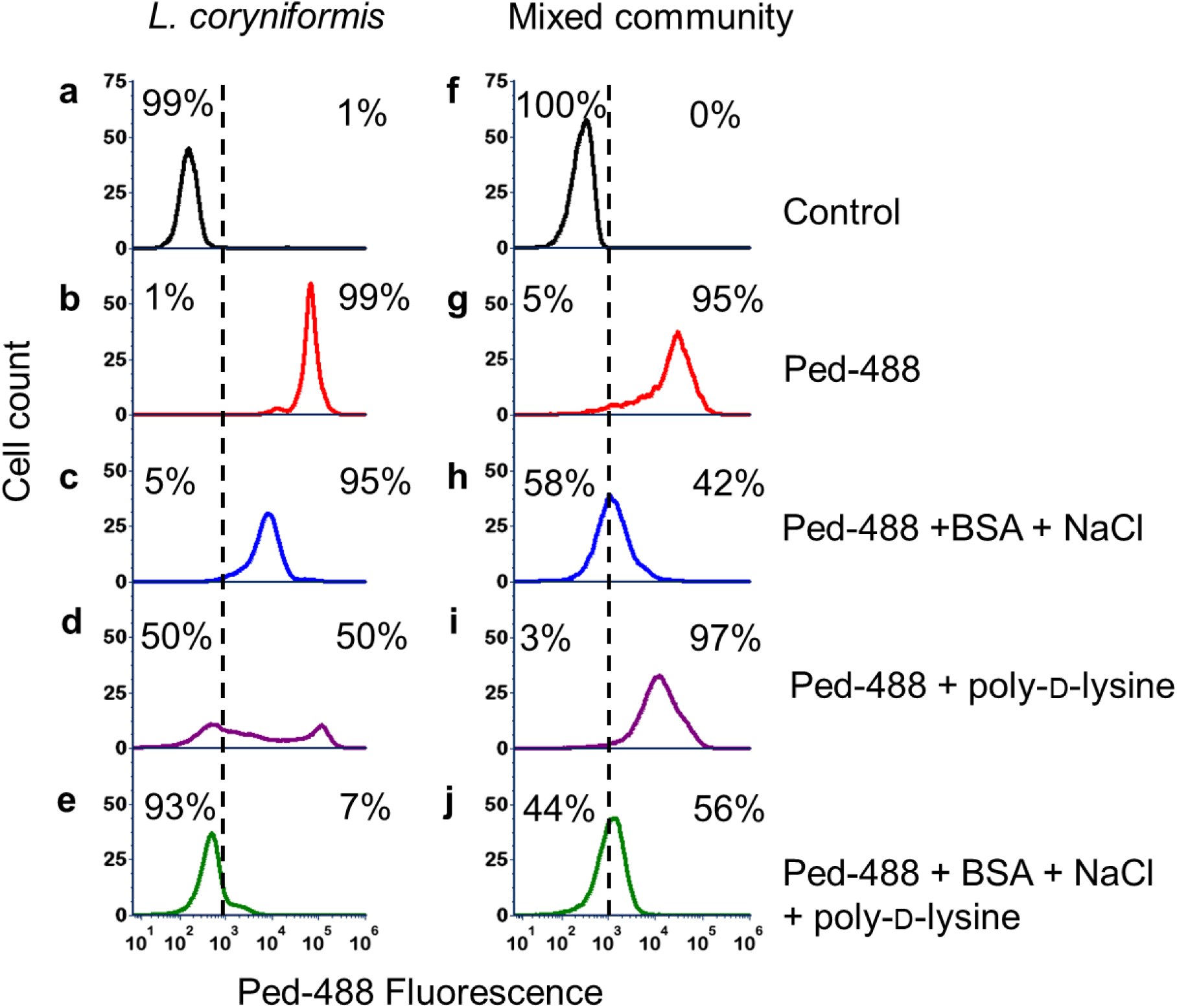
Pediocin binding in presence of different solutes and other microorganisms. Binding of pediocin PA-1 (0.25 μM) to (a−e) *L. coryniformis* (OD_600_ of 0.1), and (f−j) mixed community comprising of *B. subtilis*, *L. plantarum*, *W. confusa*, *L. casei*, *E. faecium*, *E. coli* (overall OD_600_ of 0.1) in presence of various solutes – BSA (1 %), NaCl (100 mM) and / or poly-D-lysine (PDL) (100 μM)

Recent literature attributes change in cell surface charge to emergence of microbial resistance against AMPs (4). This is usually achieved by modification of lipid A in Gram-negative and lipid II in Gram-positive organisms (38). In light of the results we are present here, it may be interesting to see if these modifications also affect the hydrophobicity, which might, in turn, affect the potency of pediocin and other AMPs to the target species. Detailed understanding of this feature may give us another handle to combat drug resistant microbes.

### Presence of other cells decreases pediocin PA-1 binding and potency to its target

Finally, we wanted to test how presence of non-target species impact the binding of pediocin PA-1 to its target, *L. coryniformis* (Fig. 6). In order to distinguish the target from non-targets (*L. casei*, *L. plantarum*, *W. confusa*, *E. faceium*, *B. subtilis* and *E. coli*), we stained *L. coryniformis* with CellBrite 640 (Fig. 6b–c). We confirmed that staining with CellBrite 640 did not affect the binding of Ped-488 to *L. coryniformis* (Fig. S5). When *L. coryniformis* stained with CellBrite 640 was exposed to Ped-488, nearly all the cells (86 %) showed green-shift indicating that pediocin was bound to the cells (Fig. 6c). However, in presence of the non-target species, only 42 % of the *L. coryniformis* cells showed binding to pediocin. Further, this indicates that cell surface of the non-target species competes for the available molecules of the pediocin. This reduced availability of pediocin may have marked impact in the bioactivity and efficacy against the pathogens.

**Figure 6.**
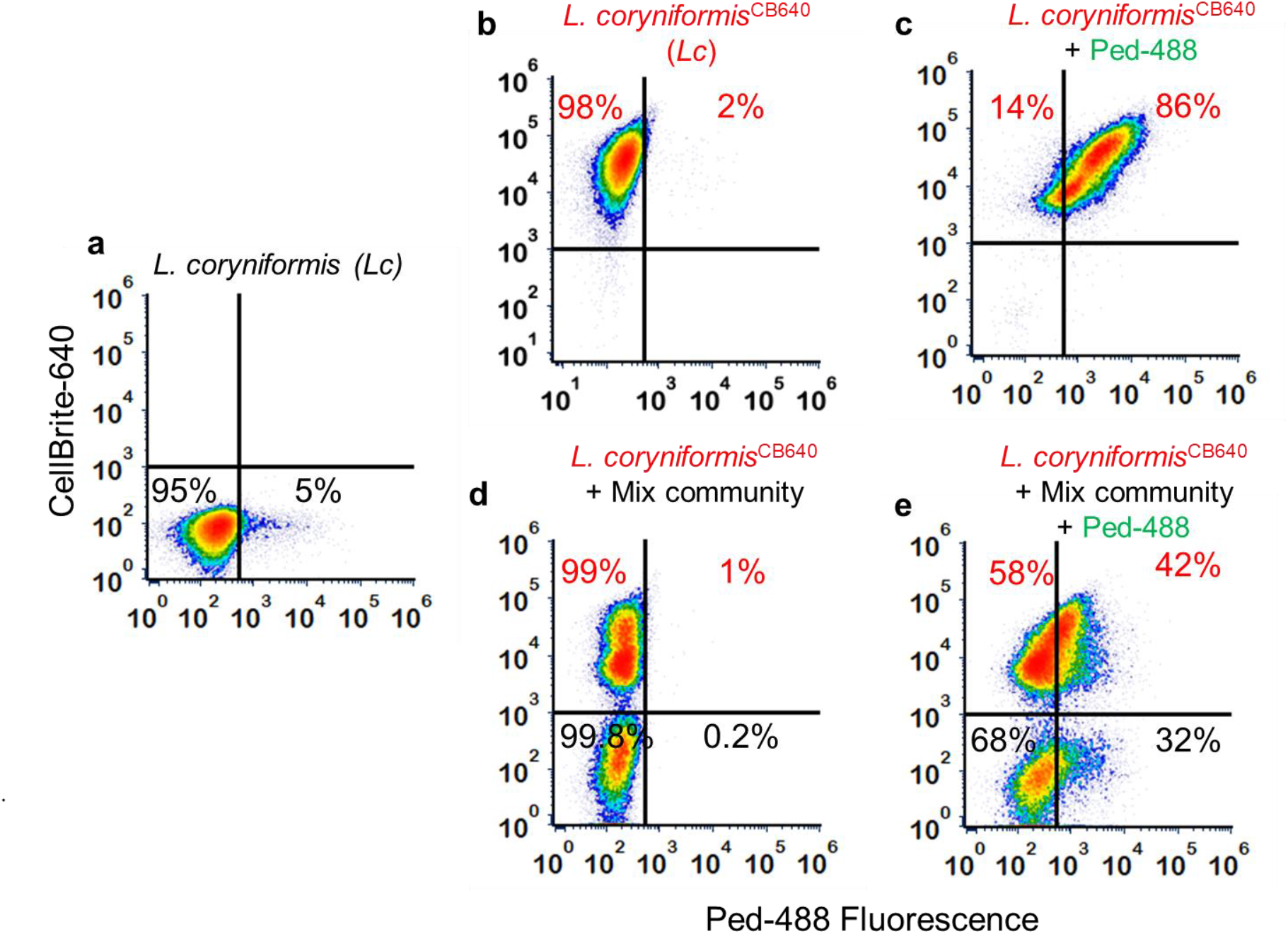
Effect of non-target species on pediocin PA-1 binding. (a) Unlabelled population of *L. coryniformis*, (b) *L. coryniformi*s labelled with CellBrite640 (CB640) to distinguish it from the non-target species, (c) *L. coryniformi*^CB640^ probed with Ped-488, (d) Equal OD mixture of *L. coryniformi*s^CB640^ and mix community, (e) Equal OD mixture of *L. coryniformi*s^CB640^ and mix community probed with Ped-488

We further tested the effect of non-target and target species on the sequestration of pediocin and the resultant decrease in potency or activity (Fig. 7). We used *L. seeliger*i (target, binder), *E. coli* (non-target, poor binder) and *B. subtilis* (non-target, strong binder) for this assay. In this pull-down assay, we incubated pediocin containing supernatant with different cell types and then measured the residual activity using agar-well diffusion assay. In control, the pediocin containing supernatant shows 1600 AU/mL (Fig. 7a) which on pre-incubation with *L. seeligeri* drops to 200 AU/mL (residual activity 12.5 %, Fig. 7b). Pre-incubation of the supernatant with *E. coli* did not affect the residual activity (1600 AU/mL, Fig. 7c). Whereas, *B. subtilis* apparently completely sequesters the pediocin from the supernatant as reflected by complete loss of activity post-incubation (0 AU/mL, Fig. 7d). This is an important observation as most of the studies have focused on effect of sequestration of AMPs by host cells or proteins on the apparent MIC *in situ* (39, 40). In recent study using a synthetic and naturally occurring peptide, ‘ARVA’ and PMAP-23, respectively, it was determined that ~107 peptide molecules/cell is required for complete killing (40, 41). However, not all peptides are affected equally by the presence of host cells like RBCs (39, 42). In the crowded environment like gut, microbial population density is much higher than the host cells (43). Due to this, it is also necessary to investigate the effect of presence of non-target species on the potency of antimicrobial peptide. In another study, using sub-MIC of LL-37 against *E. coli*, authors demonstrated emergence of non-growing and growing population. The non-growing population adsorbed the peptide, reducing free peptide concentration allowing the rest of the population to grow (44). Dead *E. coli* cells also have been shown to provide protection against LL37 (45). This is a close reflection of what would happen in natural microbial community which has a lot more diversity, density, and presence of dead cells and the components thereof. Most of these studies have been performed with human defense peptides or synthetic peptides which exhibit their affect by indiscriminate interaction with the cell membrane.

**Figure 7.**
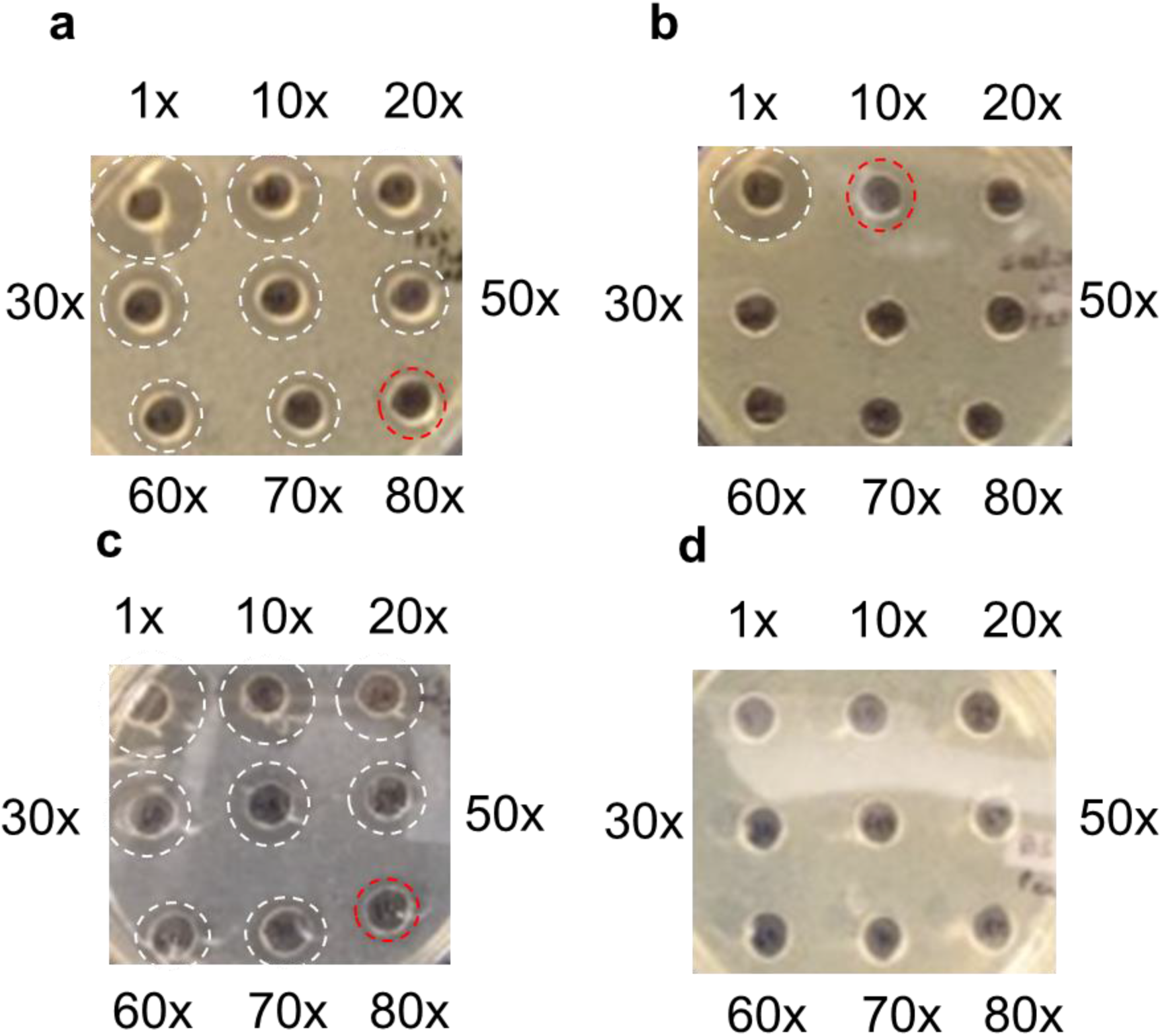
Effect of non-target species on potency of Pediocin PA-1. Agar well diffusion assay against *L. seeligeri* of (a) supernatant containing pediocin (b) pulldown supernatant after incubation with 1.0 OD_600_ of (b) *L. seeligeri*, (c) *E. coli*, (d) *B. subtilis*. White circles indicate a zone of inhibition, red circle denotes the final dilution with presence of zone of inhibition.

In conclusion, using pediocin-PA-1-fluorophore conjugate we have shown that this AMP binds to target as well as non-target species. Although pediocin PA-1 retains maximum affinity for its targets (e.g. *L. coryniformis*), other non-target species like *B. subtilis* also show significant binding to the AMP. The strength of binding is driven by the negative charge on the cell surface and is also correlated with the zeta potential. The presence of other charged solutes like salt or cationic polypeptide interferes with the binding of pediocin to target and non-target species in a more complex manner. Finally, presence of non-target species significantly reduces the availability and thus the potency of the pediocin to bind/interact with the target species. The results described here have significant implications on the use of pediocin PA-1 and similar AMPs in mixed microbial/microbiota settings, such as for use as oral antimicrobial drugs.

## Material and Methods

### Purification of pediocin PA-1

*P. acidilactici* UL5 was grown on MRS for 20 h. The culture (2L) was incubated for 30 min at 85 °C with constant stirring and pH of the culture was adjusted to 6.0 using NaOH following the color on pH-strip. The pH adjusted culture was incubated on ice for 4 h, following which cells were collected by centrifugation at 3000 ×g at 4 °C. Cells were washed thrice with ice cold sodium-phosphate buffer (5 mM, pH 6.0) and resuspended in 100 mM NaCl (100 mL). After adjusting the pH of the cell-suspension to 2.0, it was incubated at 4 °C overnight to allow the desorption of pediocin. The supernatant collected by centrifugation at 3000 ×g for 20 min, dialysed (Snakeskin® dialysis tubing, 3 kDa MWCO, ThermoFisher) against water (1:10,000 dilution) at 4 °C. Each dialysis step (1:10 dilution) was carried out for 6 h. Partially purified pediocin was lyophilized, resuspended in sodium-acetate buffer (pH 4.6, 100 mM) and stored at 4 °C. Protein was estimated using Bradford reagent (VWR).

### Bioactivity spectrum of pediocin PA-1

The antimicrobial spectrum of pediocin PA-1 was assayed using kill curve. Cultures grown overnight were resuspended in fresh medium at an OD_600_ of 0.05. EC_50_ was determined by incubating cells with varying concentration of pediocin in a 96-well plate and growth was measured at 600 nm after 24 h.

### Conjugation of pediocin PA-1

Partially-purified pediocin PA-1 was conjugated to Atto488 using 1-ethyl-3-(3-dimethylaminopropyl)carbodiimide (EDC) chemistry. Briefly, 100 μL of Atto488 (2.5 mM, dissolved in PBS), 25 μL of EDC (20 mM in PBS) and 125 μL of dimethylformamide was incubated at room temperature (RT) in dark for 10 min. Further, 28μL of *N*-hydroxysuccinimide (50 mM in PBS) was added and incubated at RT in dark for another 10 min. At this point, 250 μL of pediocin (312 μM) was mixed with the activated Atto488 and incubated at RT in dark for 2 h. The reaction was dialyzed (1:10,000) against PBS. The conjugated pediocin was then further separated from the unreacted dye using polyacrylamide desalting column (1.8 kDa MWCO). The conjugated peptide was subjected to MALDI-TOF for determination of mass and SDS-PAGE followed by visualization under UV for fluorescence signal.

### Binding assay

The binding affinity and avidity of pediocin to various Gram-positive and -negative bacteria were studied using flow-cytometry. Late log-phase cells were washed in the sodium phosphate buffer (5 mM, pH 6.0) and resuspended at an OD_600_ of 10^−3^ in sodium phosphate buffer (5 mM, pH 6.0) containing 1 % BSA and 100 mM NaCl. Cells were incubated with varying concentration of the conjugated peptide at 4 °C for 30 min. The unbound peptide was removed by centrifugation and the pellet was resuspended in sodium-phosphate buffer (5 mM, pH 6.0) containing 1 % BSA. Flow cytometry was performed on an Attune NxT flow cytometer (Life Technologies). Data (≥10^4^ events were recorded) was recorded using a green laser (488 nm) voltage of 360 V, gating was performed on SSC-A versus SSC-H plot to reduce false events.

### Zeta potential measurement

We performed zeta potential measurements to determine bacterial surface potentials. The zeta potential indicates an electrochemical property of the bacterial cell surface which represents the transmembrane potential that maintains the cell wall/membrane architecture. The zeta potential of the bacterial cell was measured in potassium-phosphate buffer (5 mM, pH 6.0) using ZetaPALS, Brookhaven. The results for zeta potential are presented as the average value from three independent cultures (6 measurements per culture).

### Binding assay in the mixed community

Late log-phase cells were washed in the sodium phosphate buffer (5 mM, pH 6.0) and resuspended at an OD_600_ of 0.1 in sodium phosphate buffer (5 mM, pH 6.0) in presence or absence of 1 % BSA and 100 mM NaCl and / or 100 μM poly-D-lysine. *L. coryniformis* was stained with CellBrite 640 (Biotium, USA) as per manufacturer’s instructions before being re-suspended in the assay conditions. Cells were mixed in equimolar ratios and incubated with 125 μM of Ped-488 at 4 °C for 30 min. The unbound peptide was removed by centrifugation and the pellet was resuspended in sodium-phosphate buffer (5 mM, pH 6.0) containing 1 % BSA. Flow cytometry was performed on an Attune NxT flow cytometer (Life Technologies). Data (≥10^5^ events were recorded) was recorded using a green (488 nm) and red laser (640 nm) gating was performed on SSC-A versus SSC-H plot to reduce false events.

## Acknowledgments

We would like to thank the Nair lab members – Todd C. Chappell and Josef R. Bober, in particular – and Prof. James A, Van Deventer (Tufts University) for help in developing protocol for cell-binding experiments. We would also like to thank Prof. Xiaocheng Jiang and Yixin Zhang (Tufts University) for allowing access, and help with, the ZetaPALS analyzer. We would also like to thank Prof. Ismail Fliss (Laval University) for sending us *Pediococcus acidilactici* UL5.

