## Supplemental Information for "Charged Gram-positive species sequester and decrease the potency of pediocin PA-1 in mixed microbial settings"

**Table S1: Strains used in the study**

| Organism | Pediocin resistance |
|---|---|
| *Lactobacillus coryniformis* B-4390 | Sensitive |
| *Listeria seeligeri* B-37019 | Sensitive |
| *Bacillus subtilis* 168 | Resistant |
| *Lactococcus lactis* MG1363 | Resistant |
| *Lactobacillus casei* B-1922 | Resistant |
| *Lactobacillus plantarum* WCSF1 | Resistant |
| *Enterococcus faecium* NRRL B-2354 | Resistant |
| *Weisella confusa* sp. | Resistant |
| *Escherichia coli* DH5α | Resistant |

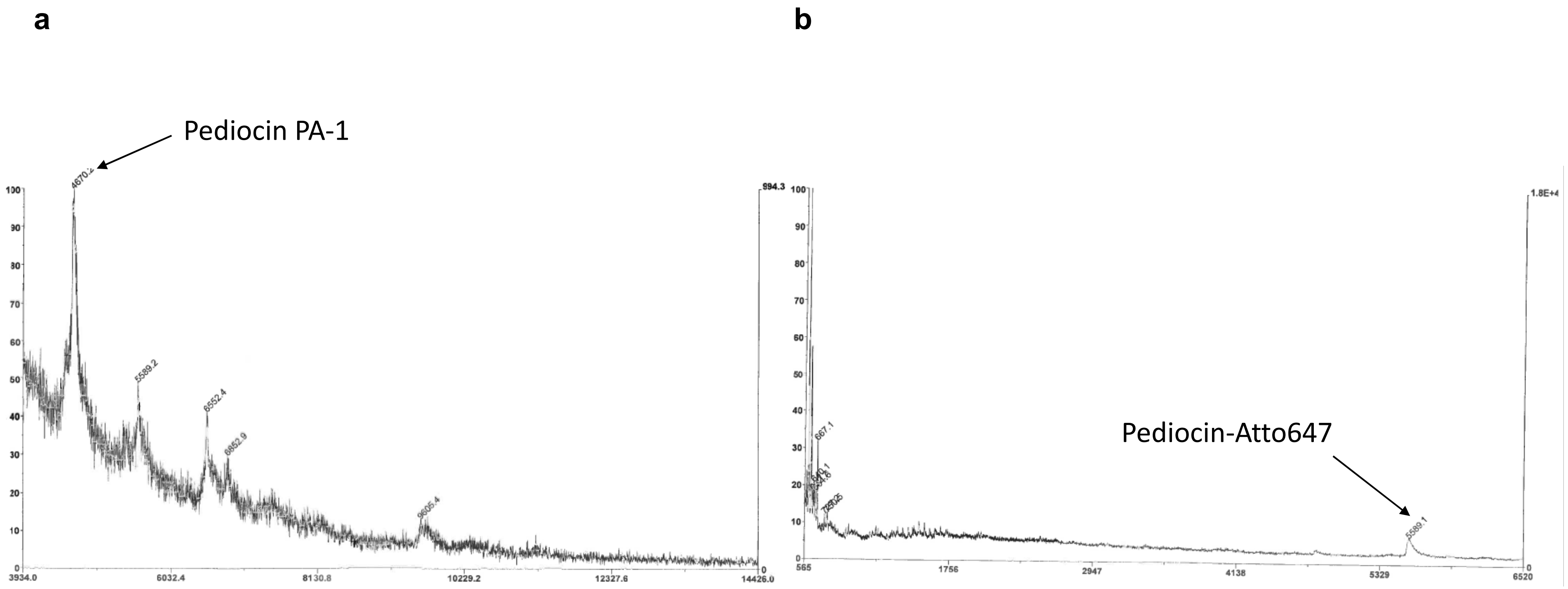


**Figure S31 MALDI-TOF-MS of (a) pediocin and (b) Ped-647**
The partially purified pediocin PA-1 gave a predominant peak at 4,670 Da, which is close to its expected molecular mass. The Atto647 conjugate gave a predominant peak at 5,589 Da, this difference in molecular mass of 919 Da is close to the molecular mass of Atto647-NHS ester (811 Da), suggesting incorporation of single molecule of dye per peptide

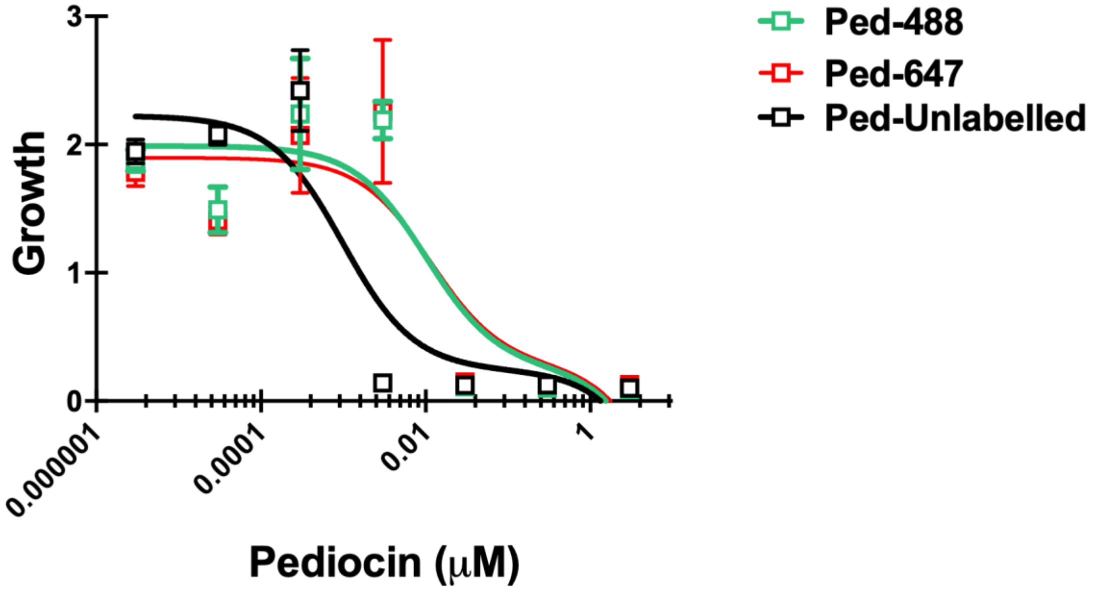


**Figure S2. Kill curve of pediocin and its conjugate against *L. coryniformis.***
The $EC_{50}$ for pediocin was determined to be 1 nM, which increased to 10 nM and 11 nM for Ped-488 and Ped-647, respectively

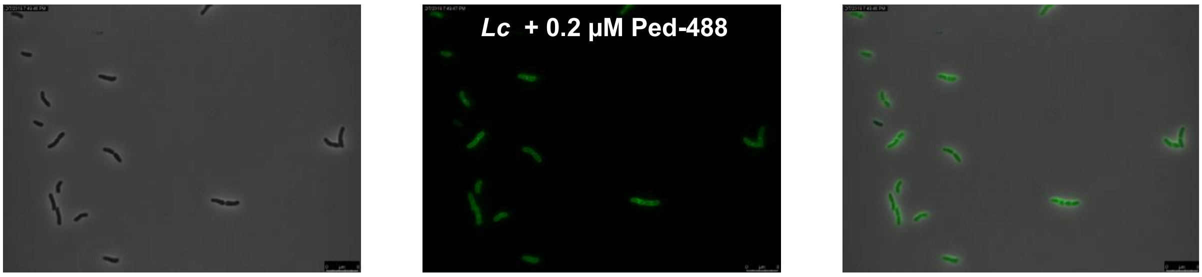


Phase GFP-filter Overlay

**Figure S3 Binding of Ped-488 to *L. coryniformis***
The stationary phase cells of *L. coryniformis* was harvested, washed with phosphate buffer (5 mM, pH 6) and incubated with 0.2 µM Ped-488 on ice before visualizing under fluorescence microscope. We could see fluorescence lining cells presumably due to binding of Ped-488 to cell membrane

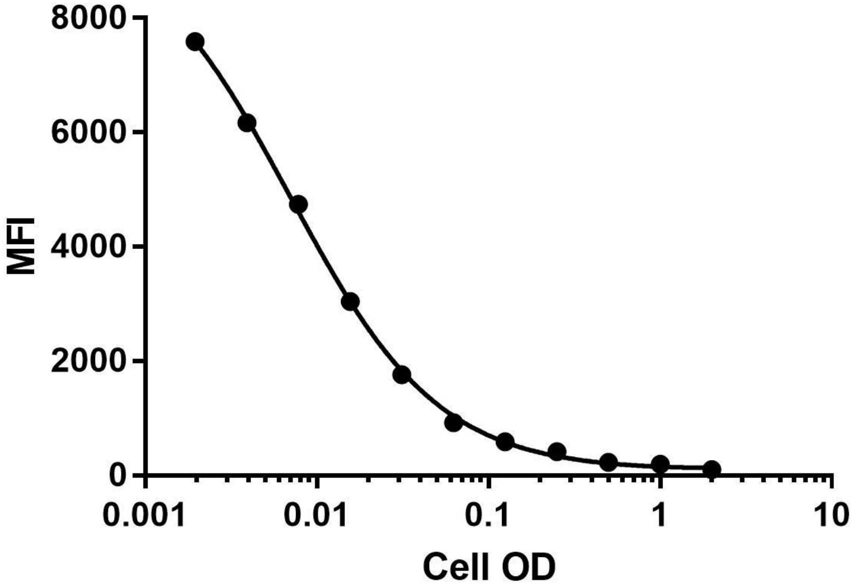


**Figure S4: Binding of Ped-488 under different cell OD**
Different cell OD of *L. coryniformis* was subjected to 100 µM of Ped-488 to determine optimal range for binding experiments. The MFI increased gradually with lower cell OD, however the △MFI between the successive cell OD dropped progressively indicating the saturation of available binding sites.

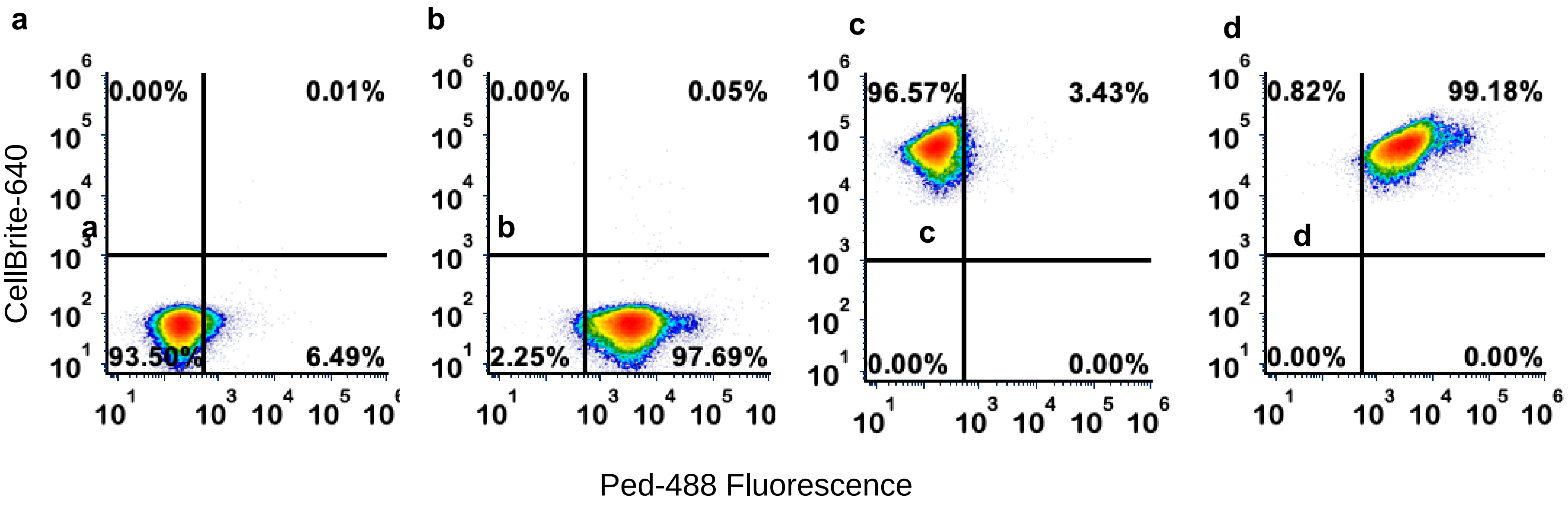


**Figure S5: Flow cytometry of Ped-488 bound *L. coryniformis***
*L. coryniformis* (a) unstained, (b) bound with Ped-488, (c) stained with CellBrite 640, (d) CellBrite 640 stained cells with Ped-488
